# Comprehensive mapping of Neurofibromin (NF1) expression in developing mouse brain

**DOI:** 10.64898/2026.03.22.713444

**Authors:** Vishal Lolam, Achira Roy

## Abstract

Neurofibromin (NF1) is a critical negative regulator of the RAS-RAF-ERK pathway, mutations in which have been clinically implicated in various neurodevelopmental disorders. However, the lack of a high-resolution spatiotemporal map has obscured the understanding of why specific cell populations and developmental processes are uniquely vulnerable to *NF1* loss. In this study, we present a comprehensive atlas of NF1 expression in the developing mouse brain. Using *in situ* hybridization and immunohistochemistry, we characterized NF1 distribution from early embryonic stages through postnatal maturation. We further integrated these findings with single-nuclei RNA-sequencing (snRNA-seq) datasets from adult mouse brain to achieve higher resolution. Our results reveal a previously undocumented graded expression pattern of NF1 across various brain regions and cell lineages. This comprehensive study will not only help understand the fundamental role of NF1 in brain development but will also be pivotal in providing a framework for studying *NF1*-associated brain disorders.

**Summary statement:** Graded expression of Neurofibromin (NF1) identified across embryonic and postnatal stages reveals unique sub-lineages of neural cells in different parts of the developing mouse brain.

## Introduction

Neurofibromatosis type 1 is one of the most common autosomal dominant neurogenetic disorders, affecting approximately 1 in 3,000 individuals worldwide (Nix et al. 2020). The condition is caused by mutations in the *NF1* gene, which encodes neurofibromin, a large multifunctional protein that serves as a critical negative regulator of the RAS-RAF-ERK pathway (Karaconji et al. 2019). NF1 consists of RAS-GTPase activating (GAP) domain, which facilitates the conversion of RAS-GTP (Active) to RAS-GDP (inactive), thereby negatively regulating RAS function (Martin et al. 1990). Since the RAS-RAF-ERK pathway is important for cell growth, proliferation and survival, NF1 acts as a brake on cellular growth and proliferation (Cichowski & Jacks 2001). In patients suffering from Neurofibromatosis type 1, this brake is lost, resulting in the formation of benign or malignant tumours in the brain and brainstem (Albers & Gutmann 2009; Lucas et al. 2022; Ayasa et al. 2024). Interestingly, a significant proportion of these pediatric patients develop neurodevelopmental disorders such as cortical malformations, epilepsy, hydrocephalus, autism spectrum disorder, intellectual disabilities, and movement disorders (Balestri et al. 2003; Sorrentino et al. 2021; Roth et al. 2019; Plasschaert et al. 2015; Hyman et al. 2005; Rietman et al. 2017; Lolam & Roy 2025; Báez-Flores et al. 2023). These clinical findings underscore the importance of NF1 in neurodevelopment beyond its canonical role in tumorigenesis.

Despite the profound impact of *NF1* mutations on the functioning of the central nervous system (CNS), our understanding of the spatiotemporal dynamics of *NF1* expression during brain development remains incomplete. Previous studies have established that NF1 is essential for embryonic development, with null mice dying in utero between embryonic days (E) 12.5 and 13.5 due to cardiovascular defects (Brannan et al. 1994; Jacks et al. 1994). In addition, studies done by knocking out *Nf1* in cultured cells and in various model systems have demonstrated altered neurodevelopmental processes (Lakkis et al. 1999; Zhu et al. 2001; Gutmann et al. 1999; Sanchez-Ortiz et al. 2014; Shin et al. 2012; Durkin et al. 2023). However, the lack of a high-resolution spatiotemporal map has limited our ability to pinpoint which developmental processes or cell populations are most vulnerable to *Nf1* loss.

Although there are some reports of NF1 expression in the brain (Daston et al. 1992; Su et al. 2007; Ryu et al. 2019; Zhu et al. 2001; Provenzano et al. 2014), no one has conducted a comprehensive high-resolution mapping of this protein in mammalian brain development. In this study, we present a high-resolution, comprehensive map of NF1 expression in the developing mouse brain. Using in situ hybridization and immunohistochemistry (IHC), we characterize the precise spatiotemporal distribution of NF1 across major brain regions and cell lineages from early embryonic stages through postnatal maturation. Moreover, we used publicly available single-nuclei RNA-seq (snRNA-seq) databases of the adult mouse brain to map NF1 expression across different cell clusters, at a resolution much higher than achievable with classical techniques alone. These results revealed a previously unreported graded expression pattern of NF1.

## Results

### NF1 is expressed in different cell lineages of the murine forebrain

Using IHC, we observed broad NF1 expression in different parts of the mouse forebrain, such as the neocortex, hippocampus, choroid plexus, thalamus, and hypothalamus (**Fig. 1A-P**). In the postnatal day (P)35 hypothalamus, NF1 strongly marked the paraventricular nucleus (PVN), suprachiasmatic nucleus (SCN), and supraoptic nucleus (SON), involved in regulating circadian rhythm, neurohormone release and fluid balance. Similar localization of *Nf1* mRNA was also observed at P35 and P0 using *in situ* hybridization (**Supplementary Fig. S1A-B”**). In embryonic ages E17.5, E14.5, and E12.5, NF1 protein expression was observed along the lateral and third ventricular linings, including neocortical and ventral telencephalic ventricular zone, cortical hem, and hippocampal primordium (**Fig. 1D-F**). Specifically, NF1 showed perinuclear expression in the neocortical plate and white matter as well as CA fields and hilus of hippocampus, but not dentate gyrus (DG) (**Fig. 1G-I**). By colabelling with different cell-type-specific telencephalic markers across development, we identified sets of cells that express the NF1 protein (**Fig. 2**). We observed strong perinuclear labelling of NF1 in excitatory pyramidal neocortical neurons from the peak neurogenic stage at E14.5 till developed cortical stage at P35 (**Fig. 2A-H,K,L**). This includes coexpression with markers specific to all mature pyramidal neurons (NeuN), deep layer-specific neurons (Ctip2), and upper layer neurons (Satb2). When checked for colocalization with the progenitors at the neocortical ventricular zone, NF1 was found perinuclearly in the Sox2^+^ apical progenitors at E17.5, E14.5, and E12.5, but not in the Tbr2^+^ intermediate progenitor cells (**Fig. 2I,J,M-P**). Further, NF1 was strongly expressed in the hippocampal CA1 and CA3 pyramidal neurons and hilar neurons but did not label any cells in the dentate gyrus (DG), as observed at P35, P0 and E17.5 (**Fig. 2Q-Y”**). These were determined by colabelling NF1 with NeuN to mark all differentiated hippocampal neurons across ages, and Ctip2, which prominently marks CA1 and developing DG at E17.5. Even at E14.5, weak NF1 expression was observed in the ventricular zone of cortical hem and hippocampal primordium/anlage (**Fig. 2Z-Z”**). E12.5 dorsal midline showed mild NF1 expression mostly concentrated along the ventricular lining (**Fig. 2AA-AA”**).

**Figure 1:**
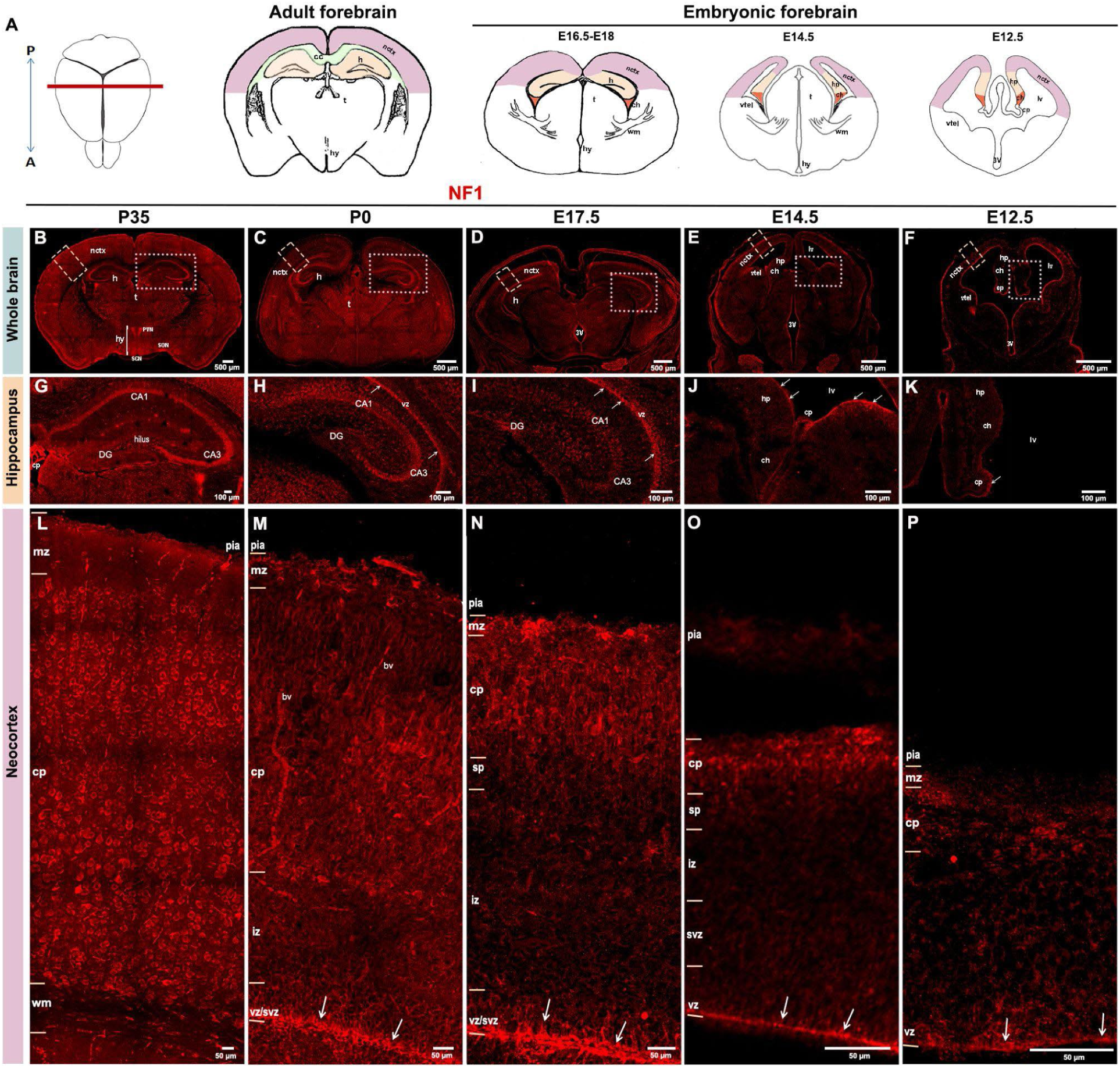
NF1 is expressed in mouse forebrain. (A) Schematics showing representative coronal sections of the mouse forebrain at adult and embryonic stages. NF1 protein expression in whole brain (B-E), hippocampus (E-I), and neocortex (J-M) across descending ages P35, P0, E17.5, E14.5, and E12.5. (B-F) NF1 was expressed in distinct zones of thalamus (t) and hypothalamus (hy) in the diencephalon. In the hypothalamus, NF1 clearly marked the paraventricular nucleus (PVN), suprachiasmatic nucleus (SCN), and supraoptic nucleus (SON) (B). Dotted boxes in (B-E) mark insets of parts of telencephalon, namely the neocortex (nctx) and hippocampus (h). (G-I) NF1 expression was observed in CA1, CA3, and hilar cells of hippocampus at E17.5, P0, and P35, but not in the dentate gyrus (DG). (J-K) NF1 was also expressed along the lateral and third ventricular linings, encompassing midline structures, namely choroid plexus (cp), cortical hem (ch), hippocampal primordium (hp), and ventral telencephalon (vtel) across ages (arrows, H-K). (L-P) NF1 is strongly expressed in all mature pyramidal neurons perinuclearly and in the ventricular/subventricular progenitor zones of neocortex across development. P, posterior; A, anterior; hp, hippocampal primordium; cc, corpus callosum; wm, white matter; 3V, third ventricle; lv, lateral ventricle; mz, marginal zone; cp, cortical plate; sp, subplate layer; iz, intermediate zone; vz, ventricular zone; svz, subventricular zone. Scalebars: 500μm (B-F); 100μm (G-K); 50μm (L-P).

**Figure 2:**
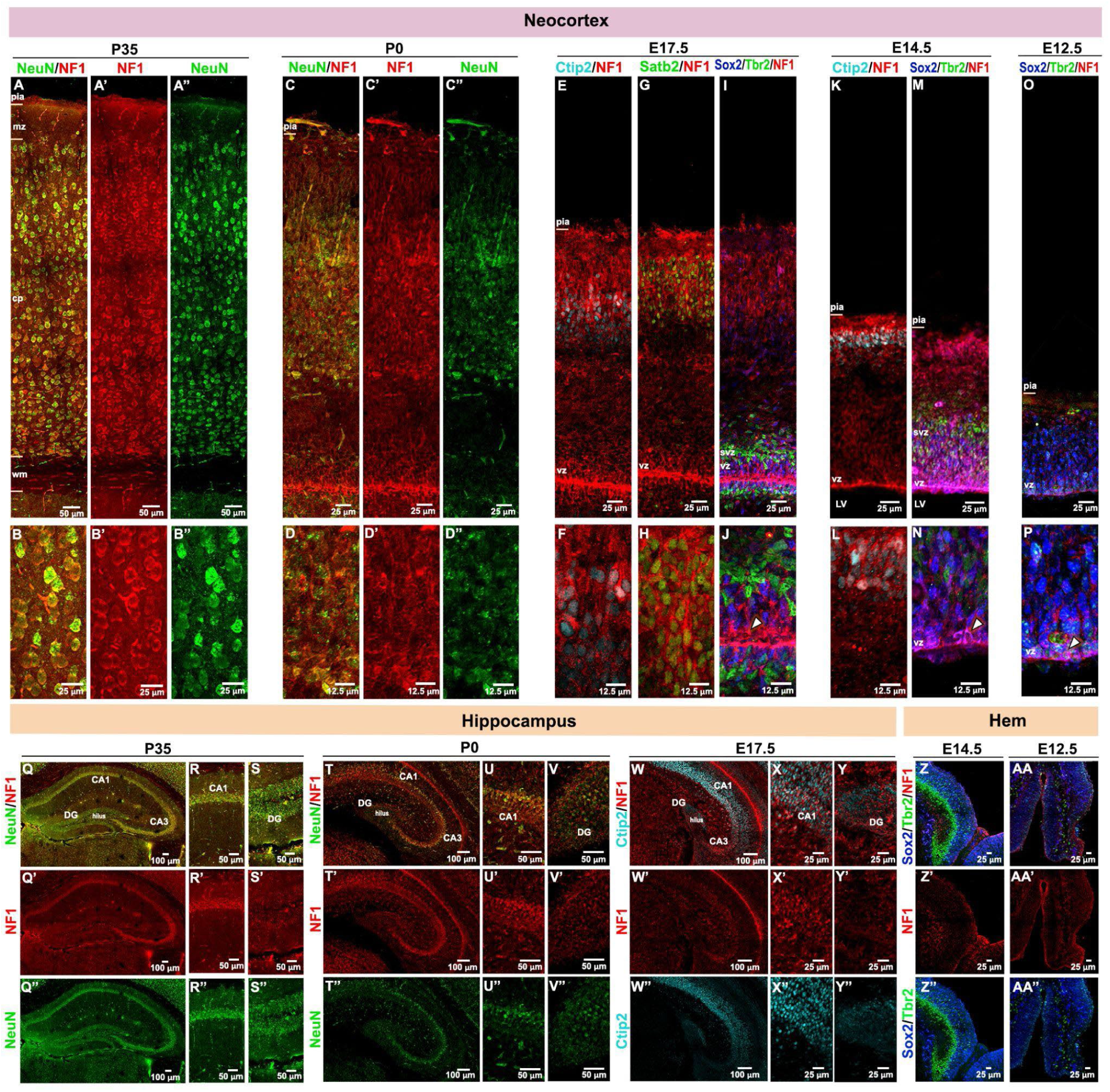
NF1 colocalizes with mature neurons and apical progenitors of forebrain. (A-D’’) NF1 colocalizes with mature neuronal marker NeuN in P35 (A-A’’) and P0 (C-C’’) neocortical plates (cp); insets show colocalizations indicated as yellow in both ages (B-B’’, D-D’’). (E-H) NF1 was expressed in both deep and upper neocortical layers. Perinuclear NF1 expression surrounded the nuclear expression of transcription factors Ctip2 and Satb2, respectively demarcating deep and upper layers. (I,J) NF1 is present perinuclearly in Sox2^+^ apical progenitors at the ventricular zone (vz) but not in Tbr2^+^ intermediate progenitor cells. (K-N) NF1 was expressed in both the developing cortical plate marked by Ctip2 and Sox2^+^ apical progenitor zone at E14.5, but not in the Tbr2^+^ subventricular zone (svz). (O,P) NF1 was expressed in the Sox2^+^ apical progenitor zone at E12.5. Arrows in J,N,P indicate perinuclear NF1 expression in the apical progenitors. (Q-Y’’) NF1 was expressed predominantly in the CA fields (CA1, CA3) and hilus of P35, P0, and E17.5 hippocampi, but no or low expression was seen in the granular neurons of dentate gyrus (DG) across ages. Hippocampal neuron-dense regions were denoted by NeuN and Ctip2 expression. (Z-AA’’) At E14.5 and E12.5, the dorsal midline of the telencephalon showed NF1 expression in choroid plexus (cp), ventricular lining, and a subset of mature neurons derived from the cortical hem (ch) and the hippocampal primordium (hp). nctx, neocortex; wm, white matter; 3V, third ventricle; LV, lateral ventricle; mz, marginal zone; cp, cortical plate; sp, subplate layer; iz, intermediate zone; vz, ventricular zone; h, hippocampus. Scalebars: 100μm (Q-Q’’, T-T’’, W-W’’); 50μm (A-A’’, R-S’’, U-V’’); 25μm (B-C”, E-I, K,M,O, X-AA’’); 12.5μm (D-D’’, F,H,J, L,N,P).

We also observed strong NF1 expression in different subtypes of cortical interneurons at P35, colabelling with Parvalbumin and Somatostatin (**Supplementary Fig. S2A-D”**). Interestingly, NF1 was observed to be present only in certain subsets of Olig2^+^ oligodendrocyte populations in the major axonal tracts of P35 mouse forebrain, namely corpus callosum, hippocampal commissure, and anterior commissure (**Fig. 3A-F’’**). This observation may suggest differential function of molecularly distinct subtypes of oligodendrocytes when NF1 levels are disrupted in the brain.

**Figure 3:**
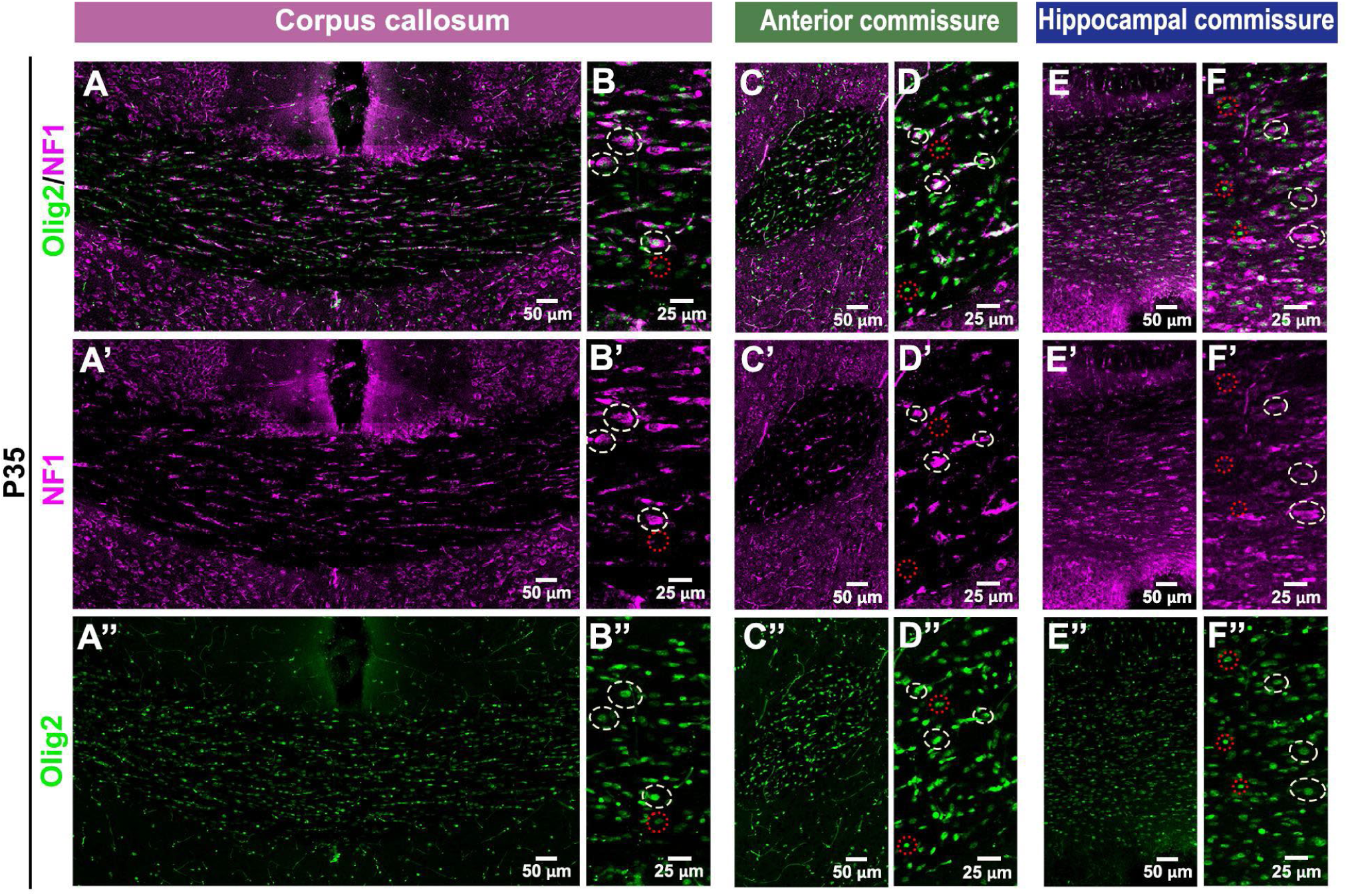
NF1 is expressed in subsets of cortical oligodendrocytes. (A-A’’) NF1 was expressed in a subset of Olig2^+^ oligodendrocytes in the main three cortical commissures of the P35 mouse brain - corpus callosum (A-B’’), anterior commissure (C-D’’), and hippocampal commissure (E-F’’). In a magnified view, cream dashed circles mark NF1^+^Olig2^+^ double-positive oligodendrocytes, while red dotted circles mark Olig2^+^ oligodendrocytes that did not express NF1 (B-B”, D-D”, F-F”). Scalebars: 50μm (A-A’’, C-C’’, E-E’’); 25μm (B-B’’, D-D’’, F-F’’).

### NF1 is expressed in different cell lineages of the murine cerebellum

Unlike the forebrain, a more restricted protein expression profile of NF1 was observed in the murine cerebellum across the developmental timeline (**Fig. 4A-J”**). Particularly, strong perinuclear NF1 expression was observed in mature and migrating Purkinje cells, colabelling with Calbindin (**Fig. 4E’,F’,G’**; **Fig. 5A-D”**). On the other hand, very low or no IHC signal of NF1 was observed at any stage of cerebellar granule cell development, when colabelled with NeuN (**Fig. 5E-H”,M,N**). *Nf1* mRNA was also observed in the parts of cerebellum at P35 and P0 using *in situ* hybridization (**Supplementary Fig. S1C-D”**). Intriguingly, NF1 expression was observed throughout the development of the cerebellar neurons, be it in the migratory state or transitory state at the embryonic nuclear transitory zone (NTZ) or later when forming the deep cerebellar nuclei (DCN) located within the white matter (**Fig. 4E”,F”,G”H”, I,I’,J,J’**). It was coexpressed with NeuN, marking neurons at the DCN, from P3 to P35 (**Fig. 5I-K”**); NeuN+ cells were not present in the prospective DCN area at P0 (**Fig. 5L-L”**).

**Figure 4:**
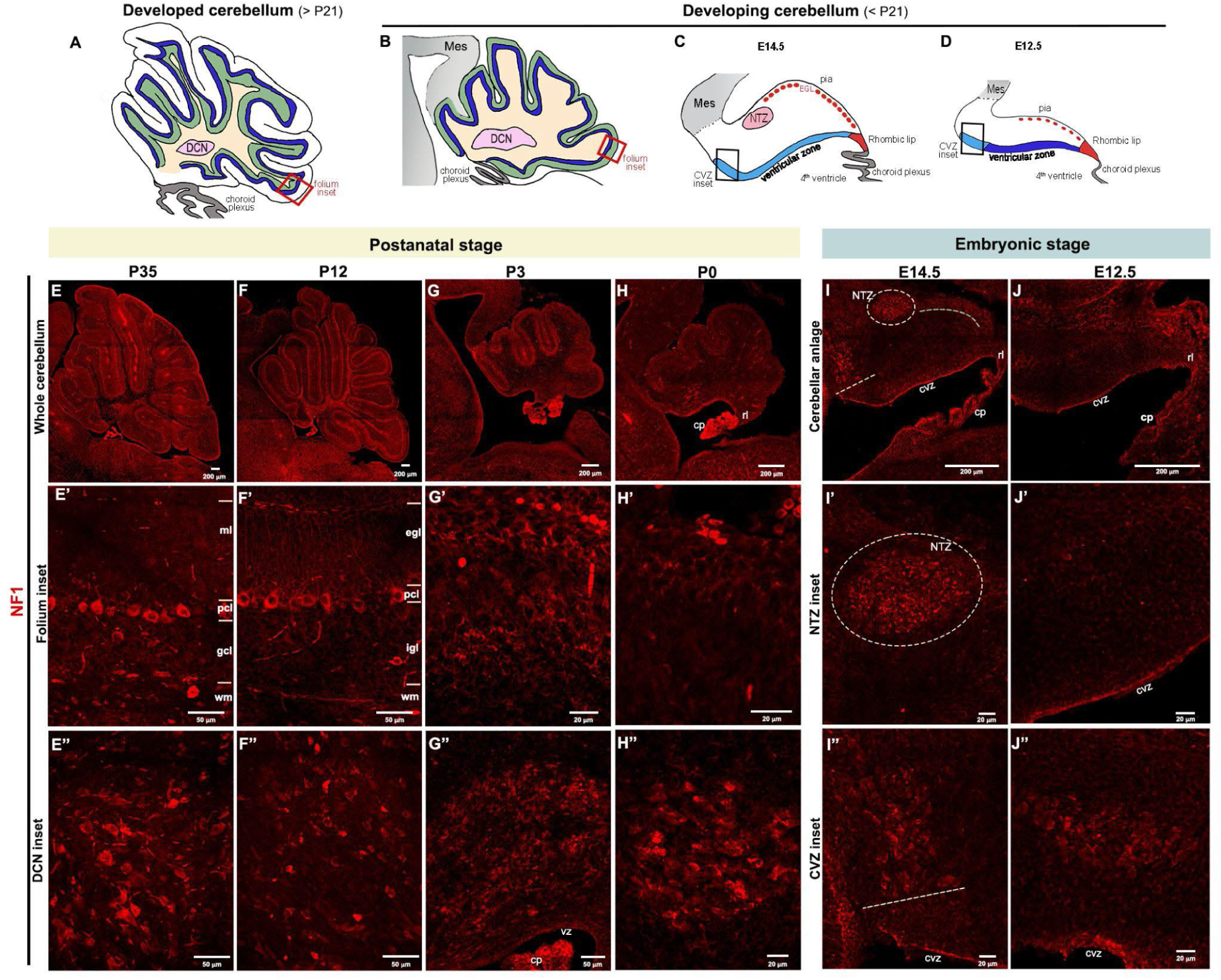
NF1 is expressed in the mouse cerebellum. (A-D) Schematics showing sagittal sections of the mouse forebrain at adult, early postnatal, and embryonic stages, demonstrating how a complex lobular laminated structure like the cerebellum develops from a simple fate-specified neuroepithelium. Boxes mark insets at P35, P12, P3 and P0 indicating magnified views of folium and cerebellar ventricular zone. (E-F’’) P35 and P12 cerebella showed strong NF1 expression in the Purkinje cell layer (pcl), cells in the white matter (wm), and in the deep cerebellar nuclei (DCN). (G-H’’) NF1 was also expressed in migrating Purkinje cells and in developing DCN at perinatal ages, P3 and P0. (I-J””) At E14.5 and E12.5, NF1 was expressed very strongly in immature cerebellar neurons of the nuclear transitory zone (NTZ), and in differentiating neurons near the cerebellar ventricular zone (cvz) and/or extra cerebellar ventricular zone. NF1 was also expressed at weaker levels in migrating rhombic lip-derived cells. NF1 was expressed in the hindbrain choroid plexus (cp) across the developmental stages, from E12.5 to the adult stage. Mes, mesencephalon; rl, rhombic lip; ml, molecular layer; gcl, granular cell layer; egl, external granular cell layer; igl, internal granular cell layer. Scalebars: 200μm (E,F,G,H,I,J); 50μm (E’-F”); 20μm (G’-H”, I’-J’’).

**Figure 5:**
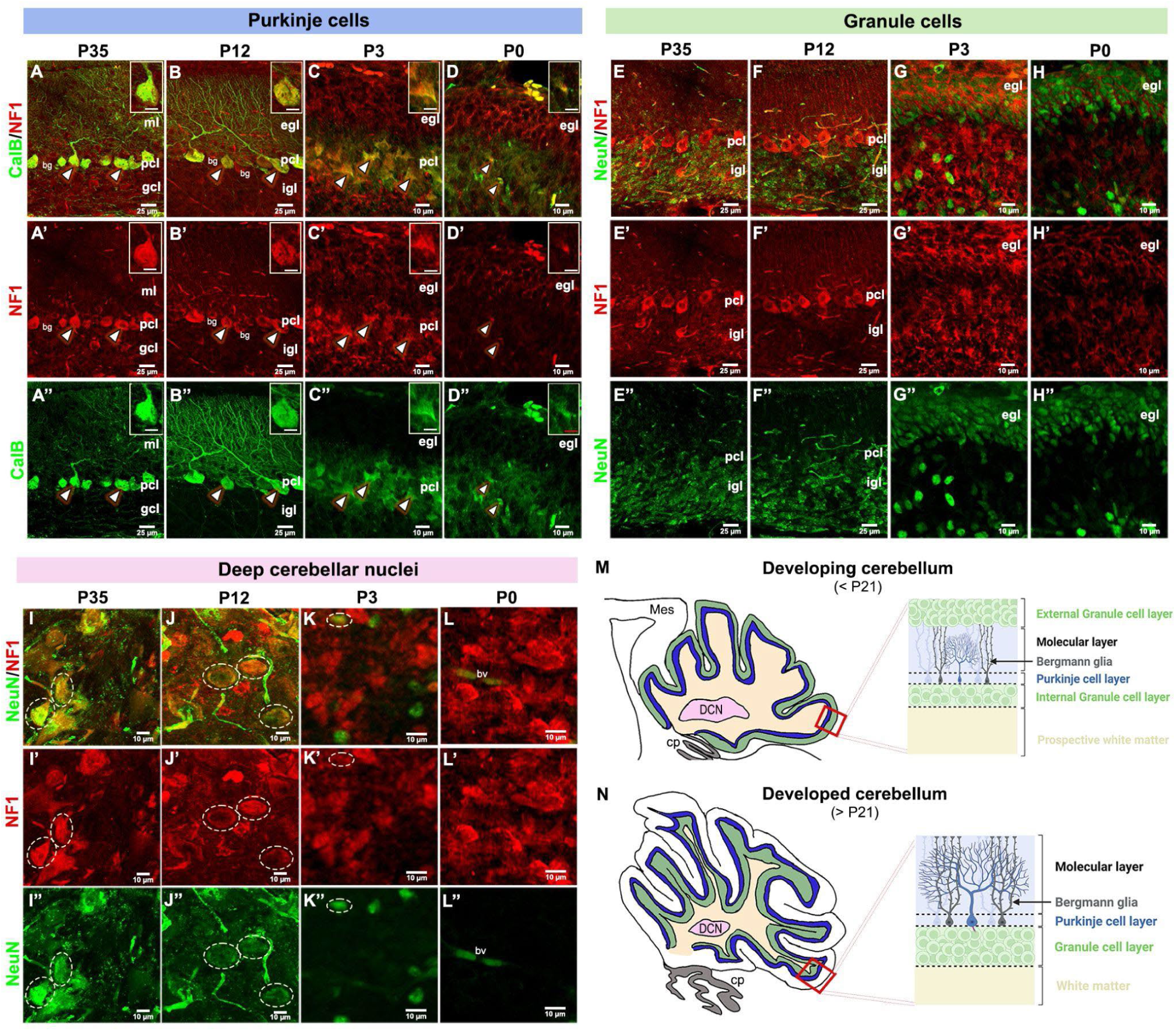
NF1 is expressed in cerebellar neuronal lineages across development. (A-D”) NF1 was expressed in Purkinje cells across development, and arrowheads show its colocalization with Calbindin (Calb). Insets show magnified views of one Calb^+^ Purkinje cell across development and its colocalization with NF1. (E-H”) Colabelling NF1 with NeuN marking cerebellar granule cells showed no or low coexpression across development. (I-L”) NF1 expression was predominant in the mature neurons of deep cerebellar nuclei (DCN) across development, colocalizing with NeuN (cream dashed circles). NeuN^+^ mature cells were not observed here at P0. (M,N) Schematics demonstrate morphology and differential lamination of developing (<P21) and developed (>P21) mouse cerebella. Mes, mesencephalon; rl, rhombic lip; ml, molecular layer; gcl, granular cell layer; egl, external granular cell layer; igl, internal granular cell layer; bv, blood vessels; cp, choroid plexus; bg, Bergmann glia. Scalebars: 25μm (A-B”, E-F”); 10μm (C-D”, G-L”).

At embryonic stages, the cerebellar anlage has distinct proliferative zones, namely rhombic lip and cerebellar ventricular zone (CVZ) (Leto et al. 2016). At early cerebellar development (E12.5), CVZ is further subdivided molecularly into the Purkinje cell-producing zone and the Pax2^+^ interneuron-producing zone. Later in development, the whole CVZ becomes potent to produce cerebellar interneurons. Perinuclear NF1 expression was observed around Sox2+ nuclei in the apical progenitors of the CVZ at E12.5 and E14.5 (**Fig. 6 A-B-B”,D,E-E”**). However, no overlap was observed in rhombic lip-derived Pax6^+^ cells at E12.5 or E14.5 (**Fig. 6C-C”, F-F”**). We also found NF1 to mark differentiated Pax2^+^ cells that have migrated out of the interneuron-producing side of the CVZ at E14.5 (**Fig. 4 I,I”, Supplementary Fig. S3A-B”**). However, NF1 was expressed in only a small subset of Tbr2+ rhombic lip-derived unipolar brush cells at E14.5 (**Supplementary Fig. S3E-E”**).

**Figure 6:**
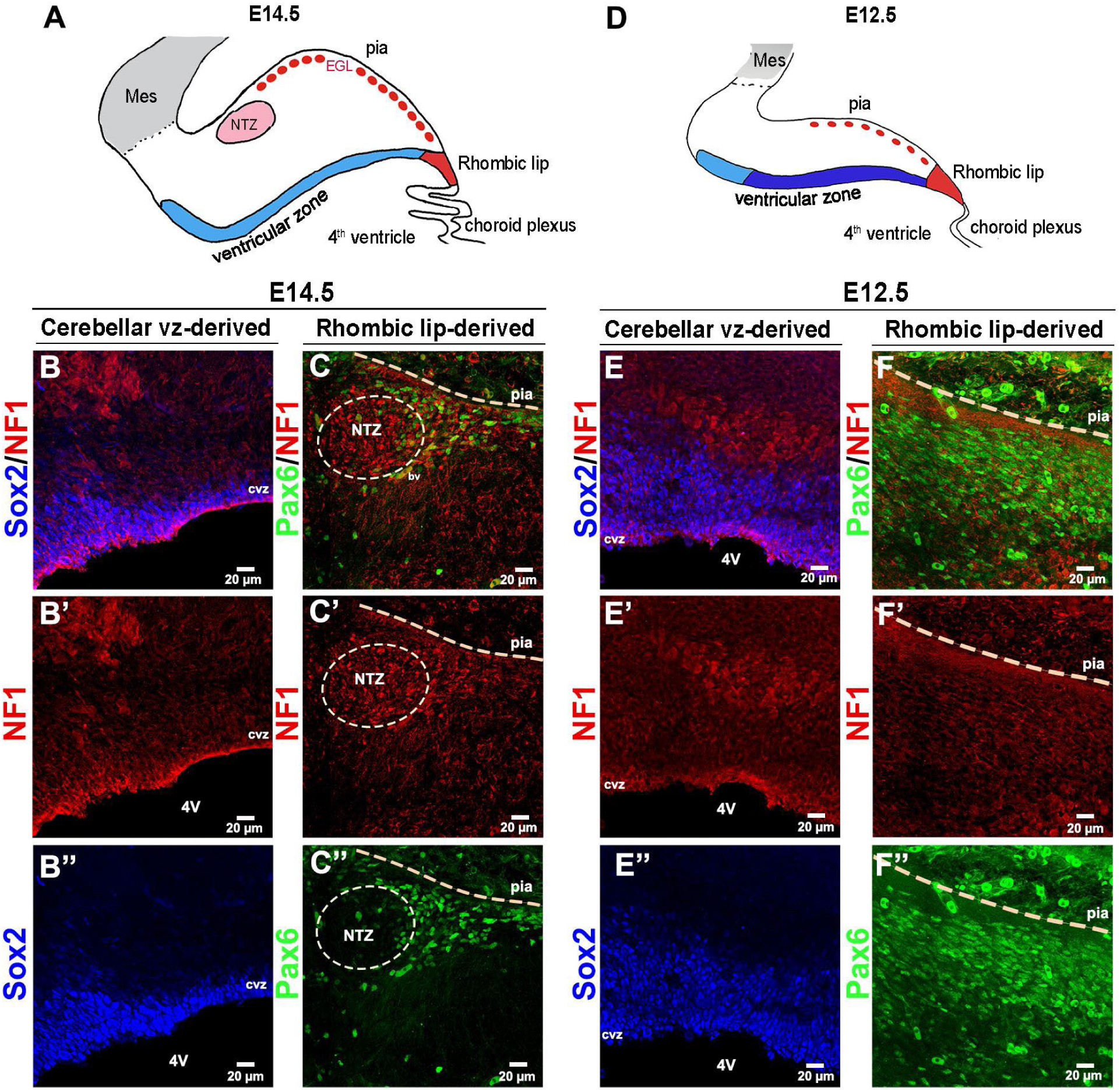
NF1 is expressed in the embryonic cerebellar ventricular progenitors. (A,D) Schematics demarcate different germinal zones of cerebellar anlage at E14.5 and E12.5. At E12.5, the cerebellar ventricular zone is subdivided into the zone producing Purkinje cells (deep blue) and the interneuron-producing zone (light blue). (B-B”, E-E”) NF1 is expressed perinuclearly in Sox2^+^ progenitors in the cerebellar ventricular zone at E12.5 and E14.5. However, NF1 was not expressed in Pax6^+^ secondary progenitor cells derived from the rhombic lip. Mes, mesencephalon; NTZ, nuclear transitory zone; 4V, fourth ventricle; cvz, cerebellar ventricular zone. Scalebars: 20μm (B-F”).

Finally, similar to forebrain results, NF1 showed an interesting expression pattern in glial lineages of the cerebellum (**Fig. 7A-N”**). At both the cerebellar folium and the DCN region, NF1 showed perinuclear expression in a subset of Olig2^+^ oligodendrocytes, as observed at P35 and P12 (**Fig. 7A-H”**). The same trend was observed in the subpopulations of Bergmann glia, which are unipolar astrocytes of the cerebellum (**Fig. 7I-N”**).

**Figure 7:**
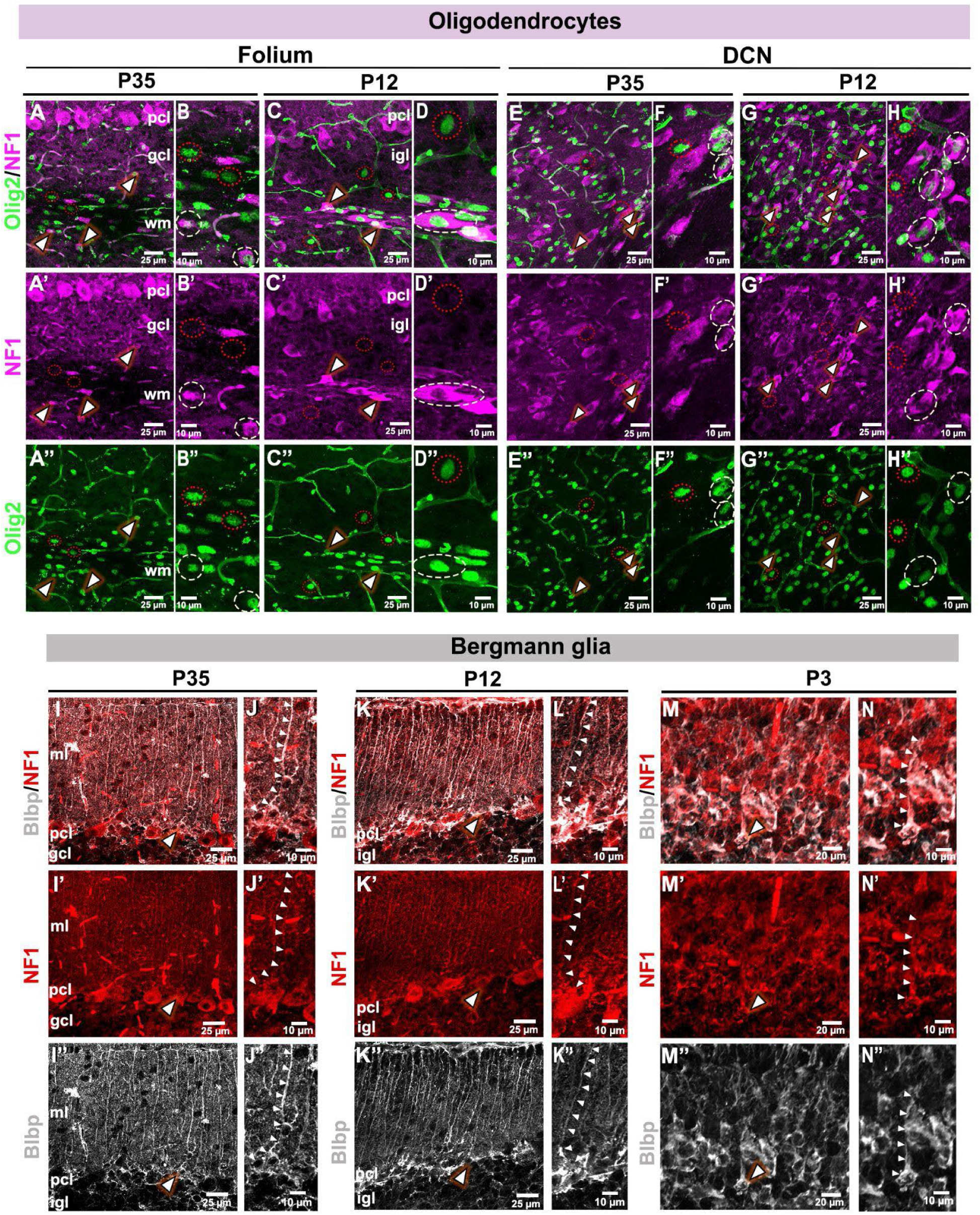
NF1 is expressed in cerebellar glial lineages. (A-H’’) NF1 was expressed in a subset of Olig2^+^ oligodendrocytes in both folium (A-D”) and deep cerebellar nuclei (E-H”) at mature (P35) and developing (P12) mouse cerebella (arrowheads). In magnified views (B-B”,D-D”,F-F”,H-H”), cream dashed circles mark NF1^+^Olig2^+^ double-positive oligodendrocytes, while red dotted circles mark Olig2^+^ oligodendrocytes that did not express NF1. (I-N”) NF1 was also expressed in the soma and projection of the Bergmann glia across development. Colocalization with Blbp is shown using arrowheads. ml, molecular layer; gcl, granular cell layer; egl, external granular cell layer; igl, internal granular cell layer; pcl, Purkinje cell layer; wm, white matter. Scalebars: 25μm (A-A”, C-C”, E-E”, G-G”, I-I”, K-K”, M-M”); 10μm (B-B”, D-D”, F-F”, H-H”, J-J”, L-L”, N-N”).

Taken together, here we report differential expression of NF1 in the mouse brain, specifically in glial populations, along development. This suggests the possibility of alternate regulatory mechanisms that eventually govern brain function.

### NF1 is expressed in distinct cell layers of the eye and olfactory bulb

Beyond regular brain regions, NF1 was also found to be expressed in accessory brain regions like the eye and olfactory bulb, both of which are connected to neurosensation (**Fig. 8A-R”**). Expression of NF1 in the developing eye was substantially strong in the lens, retinal pigmented epithelium (RPE), and neuroretina. In the neuroretina, NF1 expression was distinct in the photoreceptor cell layer (PRC), and in the retinal ganglionic cells and Müller glial cells present in the ganglionic cell layer. A subset of NF1^+^ cells in the retinal ganglionic cell (RGC) layer colocalized with Tbr2, which is known to be a key regulator of the intrinsically photosensitive RGCs (Chen et al. 2021). Sox2^+^ zone of E14.5 and E17.5 neuroretina largely did not overlap with the NF1^+^ layers, apart from the PRCs (**Fig. 8A,C-F”**). With respect to the developing olfactory bulb, NF1 was observed specifically in the mitral cell layer (MCL) of the main olfactory bulb (MOB), which are projection neurons critical for processing the sense of odours (Davison & Katz 2007). NF1 was also expressed in the accessory olfactory bulb (AOB) from P0 to P35, which is important for aggression and reproduction (Ennis & Holy 2015) (**Fig. 8G-J**). NF1 overlapped with NeuN in the MCL at P35 and P12 (**Fig. 8K,L,O-P’’**). On the other hand, it did not have much overlap with Pax6^+^ cells at P3 and P0, apart from posterior AOB (**Fig. 8M,N,Q-R”**).

**Figure 8:**
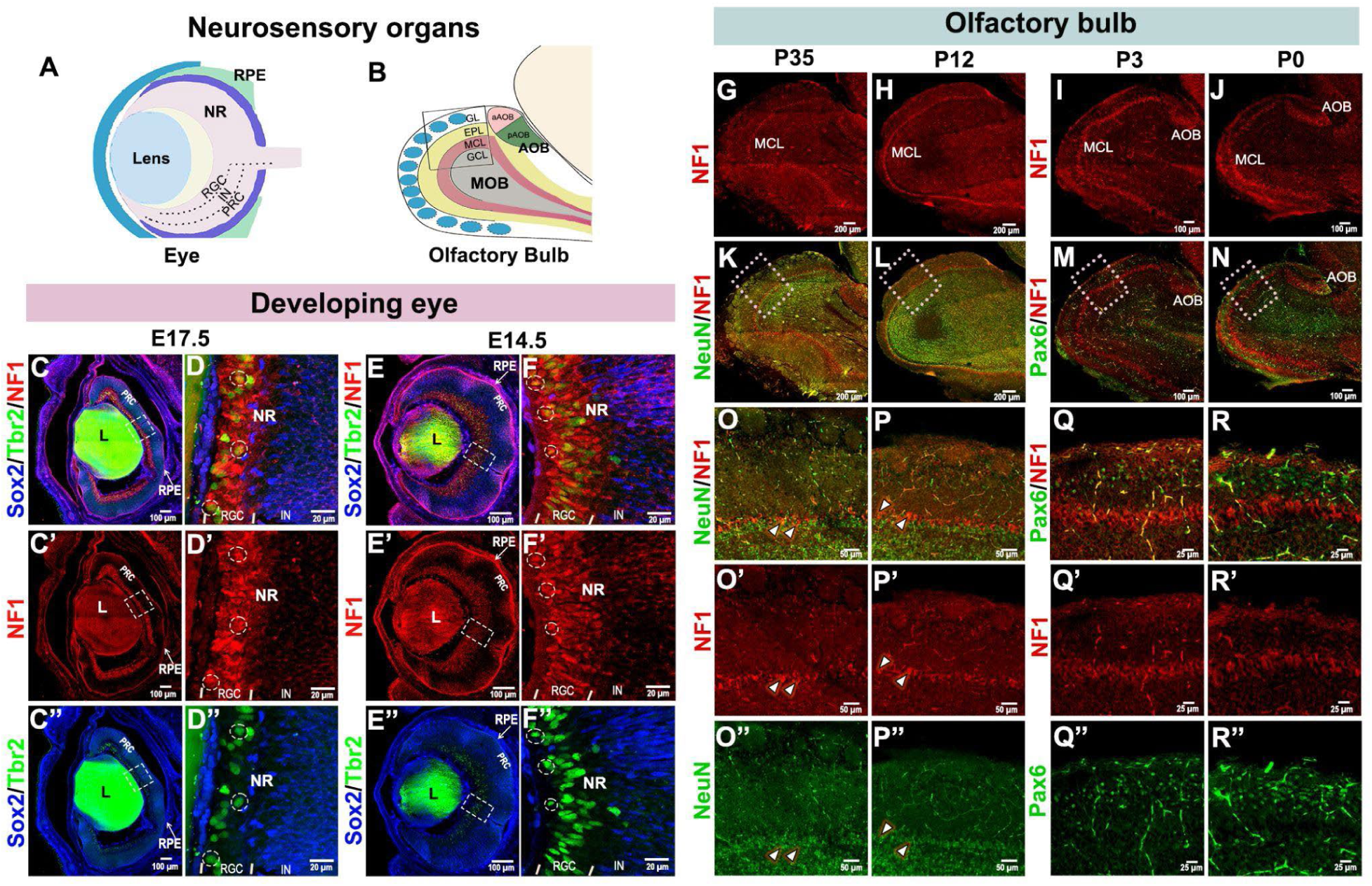
Nf1 expression in murine optic cup and olfactory bulb. (A,B) Schematics showing zones and cell layers of neurosensory organs, eye and olfactory bulb. (C,F”) NF1 is expressed in the developing mouse eye, predominantly in the lens (L) and neuroretina (NR), which includes the photoreceptor layer (PRC), retinal ganglionic cells and Müller glial cells in the broad retinal ganglionic cell layer (RGC) at E17.5 and E14.5. Magnified insets (D-D”, F-F”) show colocalization of NF1 and Tbr2 in the lens and a subset of intrinsically photosensitive RGCs. NF1 also showed wide coexpression with Sox2 in PRC and retinal pigmented epithelium (RPE), but not with the interneurons of neuroretina. NF1 colocalized with Tbr2 in the lens and a subset of (G-J) NF1 was expressed strongly across the postnatal development of the olfactory bulb. It showed strong expression in the mitral cell layer (MCL) of the main olfactory bulb (MOB) and also in the accessory olfactory bulb (AOB). (K,L,O-P”) Colabelling with NeuN at P35 and P12 identified subsets of mitral cells that were double positive for NF1 and NeuN (arrowheads). (M,N, Q-R”) Strong coexpression with Pax6 was seen in posterior AOB (pAOB) at P0 and P3, but not in the other parts of MOB. IN, interneurons; aAOB, anterior accessory olfactory bulb; EPL, external plexiform layer; GCL, granule cell layer; GL, glomerular layer. Scalebars: 200μm (G,H,K,L); 100μm (C-C”, E-E”, I,J,M,N); 50μm (O-O”, P-P”); 25μm (Q-R”); 20μm (D-D”, F-F”).

### Reanalysis of single nuclei RNAseq data reveals enriched *Nf1* expression in diverse neural cell types

To determine the expression of *Nf1* at higher resolution, we utilized publicly available single-nuclei RNAseq databases (Mortberg et al. 2023; Kozareva et al. 2021). In the adult mouse cortex, *Nf1* expression was identified in almost all the marked cortical clusters (**Fig. 9A-C**). However, detailed analysis revealed specific enrichment of *Nf1* transcripts in the lineages of cortical interneurons, oligodendrocytes, and excitatory neurons (**Fig. 9D**). In particular, *Nf1* expression was found to be highest in the oligodendrocyte lineage. Moreover, among different oligodendrocyte clusters, *Nf1* expression was found to be highest in differentiating oligodendrocytes (**Fig. 9D**). Based on our IHC data, all Olig2+ cells in the cortex were not positive for NF1. Hence, we looked at the percentage of cells positive for *Nf1* expression across different cortical clusters (**Fig. 9E**). Interestingly, oligodendrocyte precursor cells (OPCs) and differentiating OPCs showed a very high frequency of *Nf1*-expressing cells than mature oligodendrocytes. This data strongly suggests the importance of NF1 in oligodendrocyte development. Besides oligodendrocytes, cortical interneurons and excitatory pyramidal neurons also exhibit higher frequency and expression, corroborating our IHC findings.

**Figure 9:**
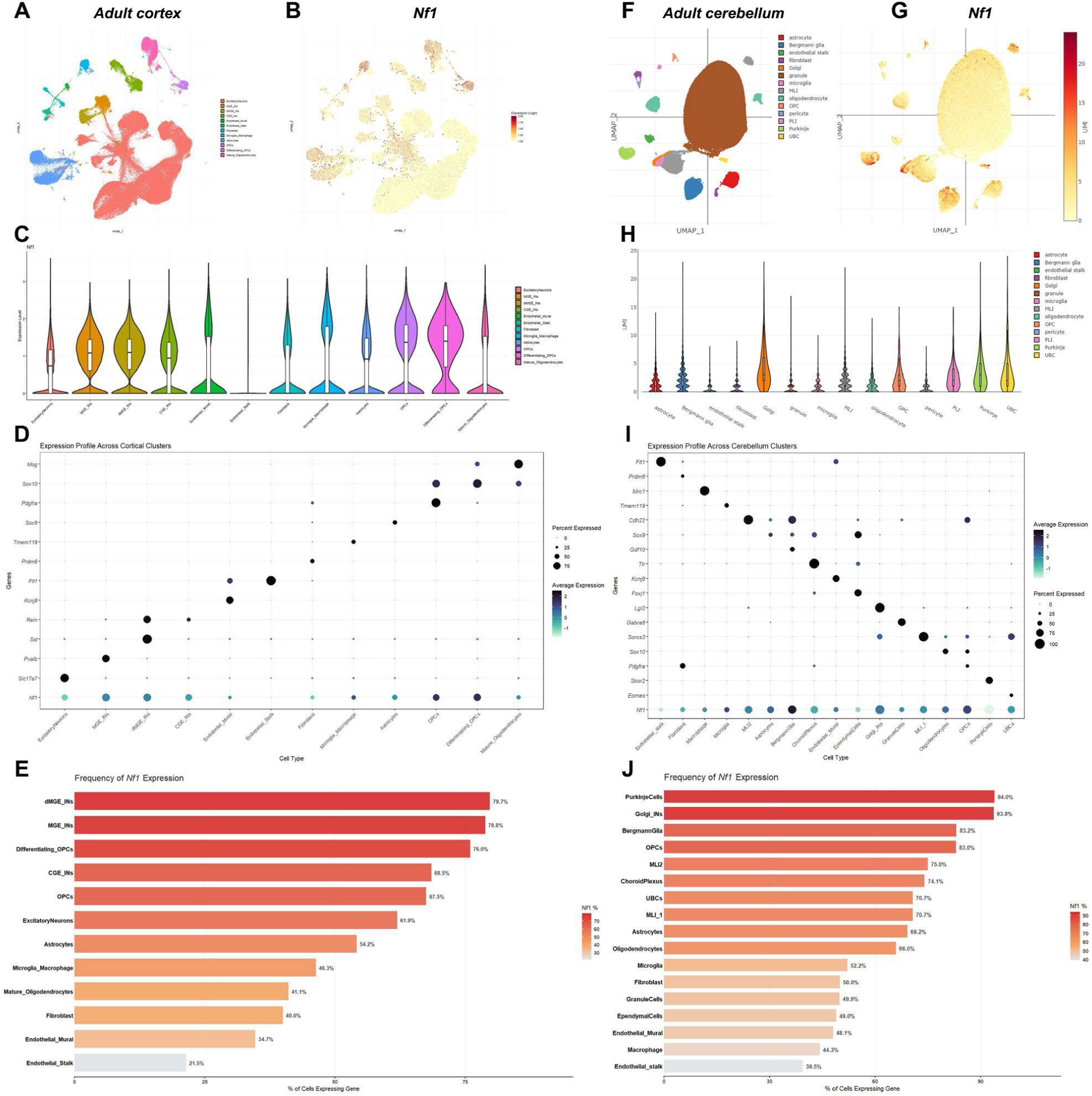
Reanalysis of single nuclei RNAseq data showing expression of *Nf1* in adult mouse brain. (A,F) UMAP plot for single nuclei RNAseq data from adult mouse cortex and cerebellum, respectively. Transcriptionally distinct clusters were coloured based on cell type. (B,G) UMAP plot illustrating the expression level of *Nf1* (red) across different cortical and cerebellar clusters, respectively. (C-E) *Nf1* expression is higher in cortical interneurons, differentiating OPCs, and excitatory neurons of adult murine cortex compared to other neural populations. (H-J) In the adult murine cerebellum, *Nf1* expression is enriched primarily in Purkinje cells, Golgi INs, Bergmann glia, and OPCs. MGE_INs, medial ganglionic eminence interneurons; dMGE_INs, dorsal medial ganglionic eminence interneurons; CGE_INs, caudal ganglionic eminence interneurons; OPCs, oligodendrocyte precursor cells; MLI, molecular layer interneurons; PLI, Purkinje layer interneurons; INs, interneurons; UBC, unipolar brush cells.

We performed a similar analysis on adult cerebellum and identified an enriched expression of *Nf1* in CVZ-derivatives, such as cerebellar interneurons, Purkinje cells, Bergmann glia, and rhombic lip-derivative unipolar brush cell (**Fig. 9F-H**). Among these clusters, the highest expression was found in the Bergmann glial cells (**Fig. 9I**). Similarly, *Nf1* showed enriched expression in the oligodendrocyte lineage of the adult neocortex, with a higher frequency of *Nf1-*expressing cells in the OPC cluster as compared to mature oligodendrocytes (**Fig. 9H**). The frequency plot also demonstrated that almost all cells in the Purkinje cell cluster were positive for *Nf1*, thus aligning with our IHC expression data.

Overall, these reanalysis results not only validated our IHC data but also helped identify *Nf1* expression in some neural cell types, such as differentiating oligodendrocytes, which are difficult to capture using conventional IHCs. Our in-depth analysis also uncovered the graded expression pattern of *Nf1* across diverse cell types in the murine adult brain. This may suggest the differential role of *Nf1* in these diverse subtypes.

## Discussion

In this comprehensive study, we demonstrate the spatiotemporal expression of neurofibromin (NF1) in the murine brain across embryonic, perinatal, and adult developmental time points, using IHC, *in situ* hybridization, and reanalyses of publicly available single-nuclei RNA-seq datasets. To our knowledge, this is the first report of an NF1 expression study, not only across major brain regions but also along almost the entire span of development. Crucially, our findings reveal molecularly distinct neural subtypes in the forebrain and cerebellum defined by the presence or absence of NF1, suggesting a previously unappreciated heterogeneity in NF1-mediated signaling during neurodevelopment. The high degree of concordance between our protein (IHC) and transcriptomic (snRNA-seq) data establishes a robust framework for understanding the role of NF1 in neural development.

### Cell-type-specific NF1 expression in the forebrain suggests roles in neuronal lineage progression and excitation-inhibition balance

In this study, we identify broad yet cell-type-specific NF1 expression across the developing and postnatal forebrain, including the neocortex, hippocampus, and major white matter tracts. NF1 demonstrated consistent perinuclear localization and strong agreement between protein and transcript levels. In the neocortex, it is expressed in excitatory pyramidal neurons across all layers and in Sox2⁺ apical progenitors, but interestingly not in Tbr2⁺ intermediate progenitors. This indicates a stage-specific role of NF1 during cortical neurogenesis. In parallel, NF1 is expressed in inhibitory interneurons, including Parvalbumin⁺ and Somatostatin⁺ populations. This suggests that NF1 is involved in both arms of cortical circuitry, raising the possibility that NF1 loss can disrupt excitation–inhibition balance. In the hippocampus, NF1 shows region-specific enrichment in the pyramidal neurons of CA fields and hilar neurons, but is notably absent from the dentate gyrus, suggesting subfield-specific functional roles. Besides, its expression is also observed in specific hypothalamic and thalamic nuclei of the diencephalon, as well as in the ventricular zone of the ventral telencephalon in younger ages. Together, these findings position NF1 as a key regulator across multiple forebrain cell types, with potential implications for lineage specification, neuronal integrity, circuit formation, and function.

### Role of NF1 in cerebellar development and motor coordination

In the cerebellum, NF1 is expressed across multiple neuronal populations throughout development, including deep cerebellar nuclei (DCN) neurons, Purkinje neurons, and cerebellar interneurons across all examined stages. In the cerebellum, proper functioning of DCN neurons and associated circuits is critical for having regular motor coordination and function (Lackey et al. 2023; van der Heijden 2024). The persistent expression of NF1 identified in the DCN and developing neurons throughout our experimental timeline highlights its importance in motor circuits. Mutations in *NF1* may impair the development or function of these neurons, resulting in motor deficits, as commonly seen in patients harbouring *NF1* mutations (Rietman et al. 2017). Similarly, defects in NF1 may affect the Purkinje cell- and cerebellar interneuron-related miscoordination. We also observed NF1 expression in subsets of Bergmann glia, cerebellar astrocytic cells that coordinate cell-cell communication, support development, and prevent neurological injury (Araujo et al. 2019; De Zeeuw & Hoogland 2015; Buffo & Rossi 2013). Having the knowledge of which subtype is affected in a disease condition may help identify specific therapeutic interventions.

### Mosaic NF1 expression across neural lineages reveals previously unrecognized cellular heterogeneity

Our analyses reveal molecular heterogeneity within both neuronal and glial populations based on NF1 expression status, identifying NF1+ve and NF1−ve subpopulations across brain regions and suggesting a mosaic pattern of expression. In particular, distinct NF1+ve and NF1−ve subsets of Olig2^+^ oligodendrocytes are observed in cerebellum and major forebrain white matter tracts, while only a subset of Bergmann glia expresses NF1. This reveals previously unappreciated diversity within glial lineages. These findings are supported both by IHC data and snRNA-seq analyses. In addition, transcriptomic data reveal NF1+ve and NF1−ve interneuron populations in the cortex and cerebellum, extending this heterogeneity to neuronal lineages. The functional significance of this differential expression remains unclear, but it suggests several possibilities: NF1 may mark distinct developmental trajectories, reflect differences in maturation state, or be dynamically regulated within lineages, defining transient cellular states. These observations raise key questions regarding the developmental origins of NF1+ve and NF1−ve populations, the mechanisms controlling NF1 expression, and the functional consequences of this heterogeneity, which may ultimately contribute to variability in NF1-associated disease phenotypes.

### NF1 expression in sensory systems extends its role to photo-dependent and olfactory circuitry

Beyond the major regions of the brain, we report NF1 expression also in specific layers of the developing eye and olfactory bulb, offering new insights into sensory-related pathologies. Specifically, NF1 is expressed in the photoreceptor layer of the neuroretina, which is vital for visual phototransduction (Hussey et al. 2022). It is also identified in Müller glial cells, which provide support and guidance for the light to reach the photoreceptor layer (Franze et al. 2007), and retinal ganglion cells (RGCs), which finally transmit sensory information to the brain. The signal perceived by the eye is known to produce both visual and non-visual responses in different parts of the diencephalon (Peirson et al. 2018). While light information cascades through the thalamic nuclei to the visual cortex for vision, it also triggers the hypothalamic nuclei critical for circadian rhythm, sleep and neuroendocrine functions (Dumbell et al. 2016; Trachtman 2010). Our study revealed NF1 expression specifically in the SCN, the master circadian rhythm generator, and in the VPN and SON, neuroendocrine nuclei involved in fluid balance and rhythmic production of hormones. Together, these strongly suggest a potential role of NF1 in light sensation, transduction, and eventually the light-dependent regulatory functions in the brain. Further work may reveal novel molecular interactions between NF1 and the diencephalon regulating sleep and metabolic homeostasis.

Similarly, the presence of NF1 expression in the mitral cells of the olfactory bulb and in both parts of the AOB suggests an underexplored role for NF1 in odor and pheromone processing and in pheromone-driven behaviors, such as reproductive and defensive responses. The conservation of these patterns suggests that NF1 may be integral to evolutionary ancient sensory circuits.

### Implications for NF1-associated neurodevelopmental disorders and tumorigenesis

This comprehensive atlas of NF1 expression across murine brain development provides a framework for linking cellular context to NF1-associated neurodevelopmental disorders and tumorigenesis. The widespread yet heterogeneous expression of NF1 across neuronal and glial populations suggests that its dysregulation may impact multiple circuits and developmental processes, contributing to the clinical variability observed in NF1 patients. The identification of NF1+ve and NF1−ve subpopulations further raises the possibility of selective cellular vulnerability, which may influence both disease manifestation and tumor initiation. These findings offer a basis for identifying cells of origin and critical developmental windows in NF1-associated pathologies.

Overall, this work provides a resource for understanding NF1 function in the developing brain and supports future efforts toward cell-type-specific and targeted therapeutic strategies.

### Limitations of the study

While this study provides a comprehensive spatiotemporal map of NF1 expression, several limitations should be considered. First, our analyses are primarily descriptive and based on IHCs, *in situ* hybridization, and reanalysis of existing snRNAseq datasets; therefore, they do not directly address the functional consequences of NF1 loss in specific cell types. Second, although strong concordance is observed across modalities, differences in sensitivity and resolution between protein- and transcript-level measurements may influence the detection of low-abundance or transient expression of NF1 in different cellular lineages. In addition, our conclusions are drawn from murine models, and although broadly informative, species-specific differences may limit direct translation to human biology. The observed NF1+ve and NF1−ve heterogeneity across cell populations also remains unresolved, particularly with respect to developmental origin, temporal dynamics, and regulatory mechanisms. Addressing these limitations through functional perturbation studies, higher-resolution temporal analyses, and cross-species validation will be important for further elucidating the role of NF1 in brain development and disease.

## Materials and methods

### Key resources table

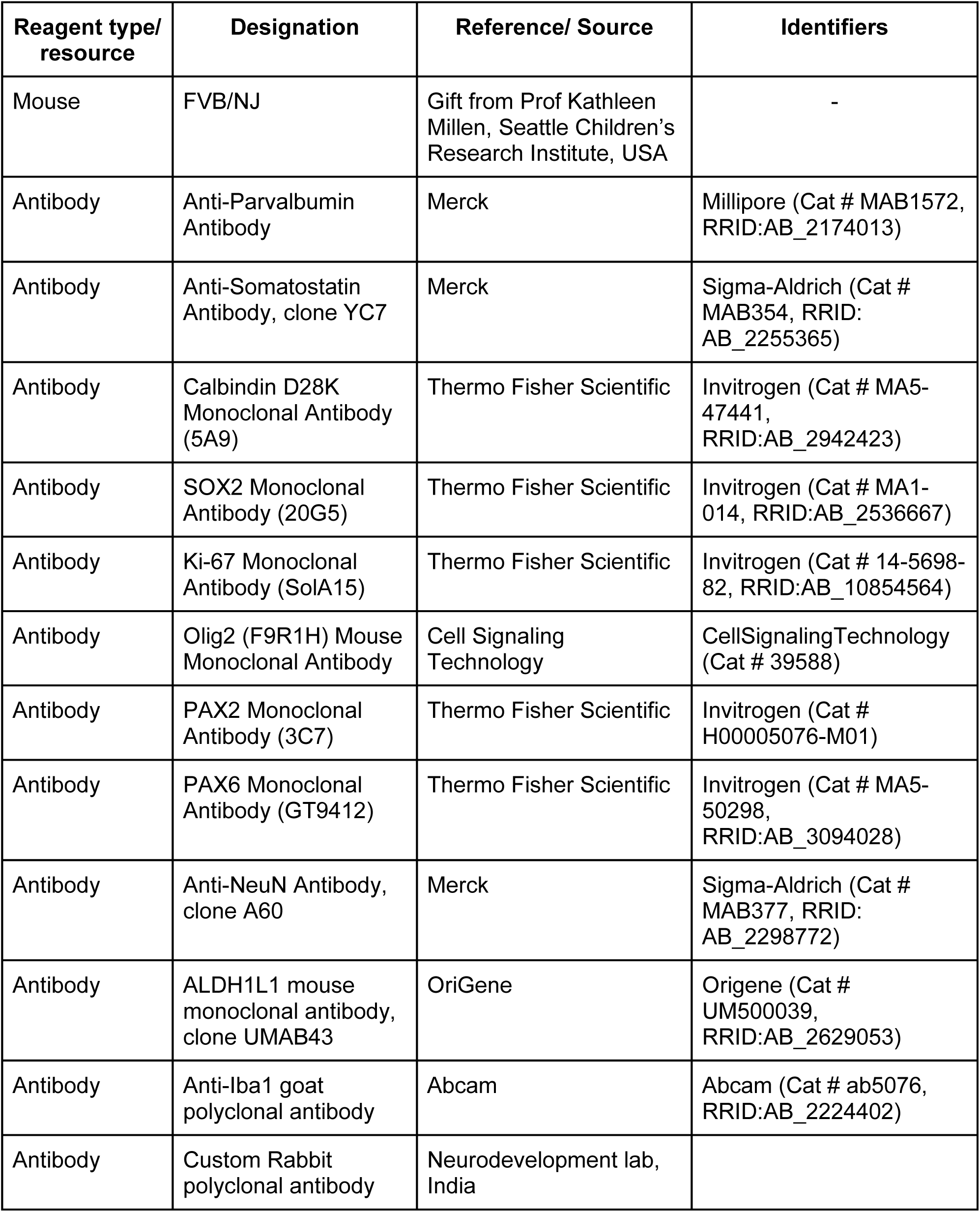
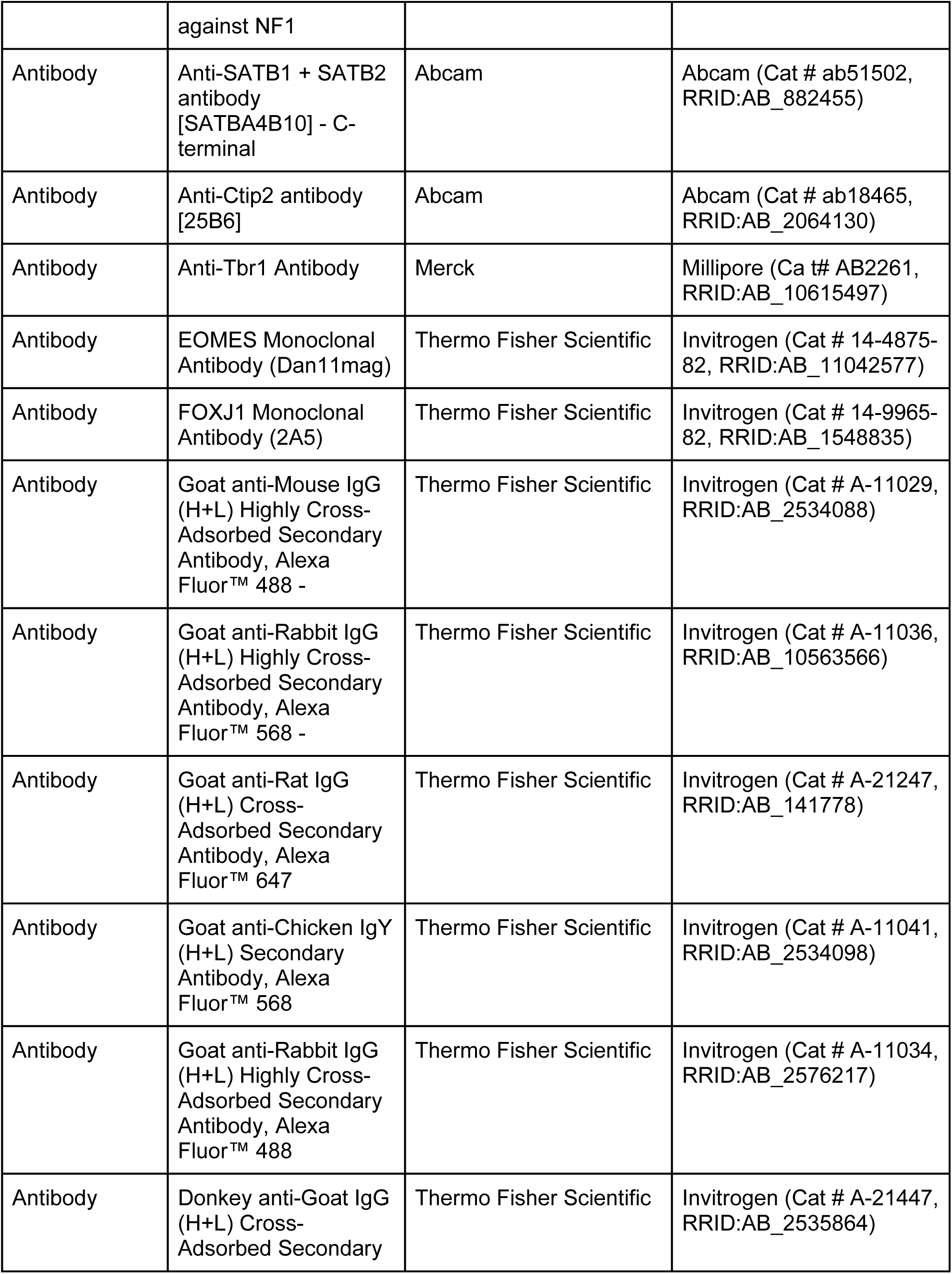
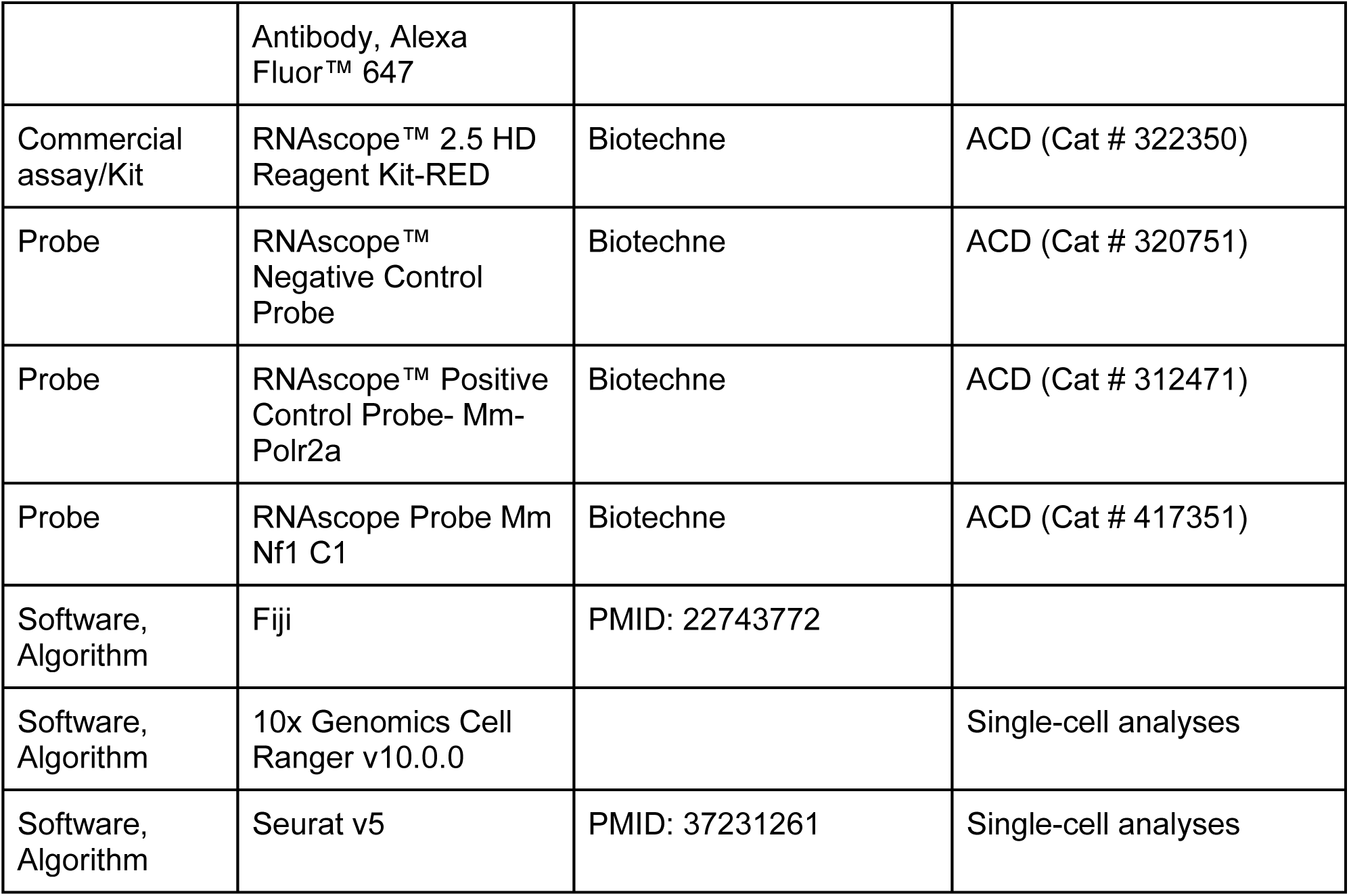

### Mice

For all the experiments FVB/NjJ mouse strain was used. Mice were housed and bred under pathogen-free conditions in the animal facility at Jawaharlal Nehru Centre for Advanced Scientific Research (JNCASR, India). All mice received food and water *ad libitum* and were subjected to a 12-hour light-dark cycle. Noon of the day of vaginal plug was designated as embryonic day 0.5 (E0.5). The day of birth was designated as postnatal day 0 (P0), which usually occurs between E19-20.

### Sample preparation

Brains from postnatal pups were harvested in phosphate buffer saline (PBS), then were fixed in 10% Formalin Solution neutral buffer (10% NBF) for 48-72 hrs for P35 and 2-6hrs for all embryonic and early postnatal ages, equilibrated in 30% (wt/vol) sucrose made in 1X PBS, and sectioned at 30 µm on a freezing microtome (Leica, Germany). These sections were then processed for *in situ* hybridization or immunohistochemical procedures.

### Immunohistochemistry

boiled in 10mM sodium citrate solution for antigen retrieval, blocked in 10% serum in PBS with 0.1% Triton X-100, and then incubated for 36-48hrs at 4°C with primary antibodies. The next day, sections were washed thrice in PBS, incubated with appropriate species-specific secondary antibodies conjugated with Alexa 488, 568, or 647 fluorophores (Invitrogen) for 2 hrs at room temperature and then counterstained with 4′,6-diamidino-2-phenylindole (DAPI) to visualize the nuclei. Sections were coverslipped using Fluoromount G mounting medium (Fisher Scientific, USA).

Information regarding the antibodies used in this study is listed in the Key Resources table above. Immunohistochemistry experiments were replicated with biologically independent samples (minimum three). All attempts for replication were successful. No outliers were encountered. Each antibody was validated for mouse and application (IHC) by the respective manufacturer, and the data are publicly available. This was also validated by us in our experiments, replicating published/expected expressions in control tissue.

### *In situ* hybridization

Assays were run using commercially available probes from Advanced Cell Diagnostics (ACD, Biotechne). Manufacturer-recommended protocols were used without modification. Sections were probed using *Nf1* (Cat# 417351) and counterstained with hematoxylin.

### Imaging of brain sections

Imaging of IHC brain sections was done using CellVoyager™ CQ1 Benchtop High-Content Analysis System using CellPathfinder (Yokogawa, Japan), and later processed in ImageJ software (NIH, Bethesda, Maryland, USA). *In situ* slides were visualized under an Olympus VS 200 slide scanner brightfield microscope. Apart from minor adjustments to contrast and brightness across the entire image, there was no further image alteration. Pseudo-colouring was used to highlight the cells of interest. Figures were prepared on Adobe Illustrator.

### Reanalysis of single-nuclei transcriptomics datasets

Using Single Cell Portal (singlecell.broadinstitute.org/single_cell), we conducted a reanalysis of single-nuclei RNAseq data of adult cerebellum (SCP795) and adult cortex (SCP2126), originally reported by (Mortberg et al. 2023) and (Kozareva et al. 2021), respectively. The Seurat object was downloaded from the portal, and downstream analysis was performed using Seurat v5 (Hao et al. 2024). UMAP clusters were annotated to different neural cell types based on their transcriptomic signatures. We specifically focused on examining Nf1 expression levels across diverse cell types.

Subsequently, we generated graphical plots with the help of ggplot2 package (Wickham 2016).

## Author Contributions

VL and AR contributed to the study conception, study design, data acquisition, data analysis, and interpretation. VL and AR wrote the first draft of the manuscript, reviewed it, and approved the final version. AR provided funding and infrastructural resources.

## Acknowledgements

We thank Prof Kathleen J Millen, Seattle Children’s Research Institute, for gifting the FVB mouse line; Varun M. and Shruti Srivastava for technical support during image acquisition. Cerebellar lamination schematics in Fig. 5M and 5N have been created using the BioRender software (https://www.biorender.com/).

## Ethics

Animal experimentation: All animal experimentation was conducted in accordance with the guidelines laid down by the Institutional Animal Ethics Committees (IAEC) of Jawaharlal Nehru Centre for Advanced Scientific Research, India (protocols AR 002 and AR 003).

## Data/resource availability

All relevant data and details on resources are provided in the article and its supplementary information.

## Diversity and inclusion statement

Preparation of this document has involved diversity and inclusion practices relevant to both the scientific content of the paper and authorship and attribution.

