## Supplementary File for "Comprehensive mapping of Neurofibromin (NF1) expression in developing mouse brain"

***Research article***

Assistant Professor

Neurodevelopment Lab, Room: SBS-B201, Neuroscience Unit,

Jawaharlal Nehru Centre for Advanced Scientific Research (JNCASR),

Rachenahalli Lake Road, Jakkur PO, Bangalore, India. PIN: 560064

**Short Title:** NF1 expression in mouse brain

**Keywords:** NF1, antibody, brain expression, cerebellum, single-cell reanalysis

**Supplementary Figures: 3**

**Supplementary information**

**Supplementary Figure S1: *In situ* hybridization shows *Nf1* mRNA expression in major parts of a mouse brain**

(A-B’’) mRNA expression of *Nf1* in the developing forebrain, counterstained by hematoxylin (H). Dotted boxes in (A,B) mark insets of parts of the telencephalon, namely the neocortex and hippocampus. (A’, A’’, B’, B’’) Magnified insets show higher expression of *Nf1* in hippocampal CA fields and neocortex, but no expression in dentate gyrus (DG). (C-D’’) Expression of *Nf1* mRNA in developing cerebellum. Dotted boxes in (C,D) mark insets of parts of the folium. *Nf1* expression is predominantly observed in mature (C’) as well as developing (D’) Purkinje cells. (D’) In addition, *Nf1* shows a strong signal in deep cerebellar nuclei from very early stage. Scalebars: 1mm (A,C); 500μm (B,D); 200μm (A’, B’, D’); 100μm (A”, B”,C’).

**Supplementary Figure S2: Nf1 is expressed in cortical interneurons**

(A-D”) NF1 showed strong coexpression with interneuron markers Parvalbumin (PV) and Somatostatin (SST) in the P35 cortical plate. Dashed circles in the magnified views confirmed colocalization (B-B”, D-D”). Scalebars: 50μm (A-A”, C-C”); 25μm (B-B”, D-D”).

**Supplementary Figure S3: Nf1 is expressed in cerebellar interneurons and unipolar brush cells**

(A-D”) Pax2 marks interneurons and progenitors in the cerebellar ventricular zone (CVZ) that give rise to interneurons. Perinuclear expression of NF1 was observed in Pax2+ interneurons that had migrated out of the CVZ in E14.5 and P12 (arrowheads). Magnified views show the subcellular localization of the NF1^+^Pax2^+^ interneurons in both ages (B-B”, D-D”). (E-E”) At E14.5, NF1 was expressed in a subset of Tbr2^+^ unipolar brush cells, as marked by dashed circles. Scalebars: 20μm (A-A”, C-C”, E-E”); Scalebars: 10μm (B-B”, D-D”)**.**
