## Supplementary figures and images for "Comprehensive mapping of Neurofibromin (NF1) expression in developing mouse brain"

### Supplementary Figure S1

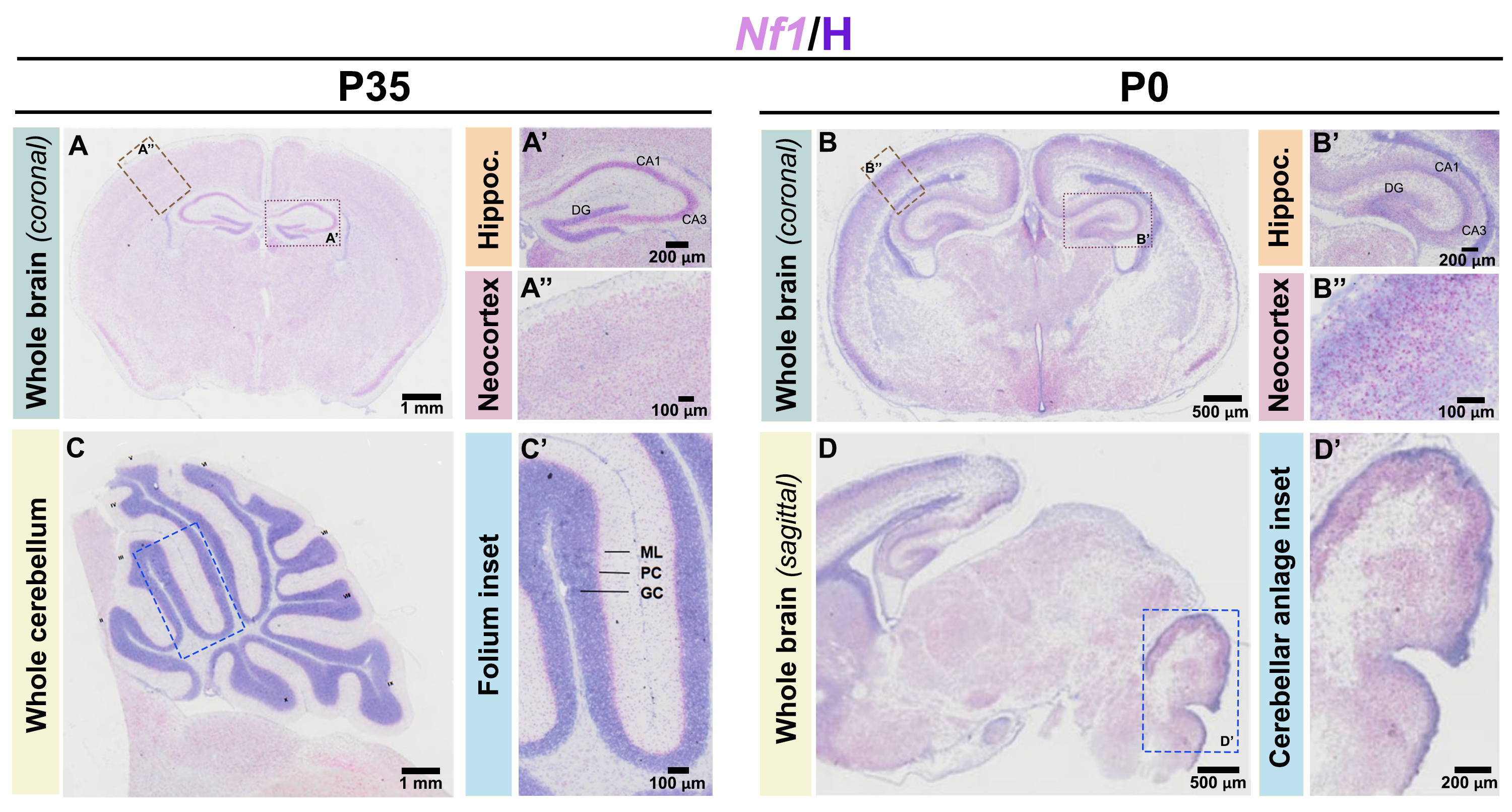

### Supplementary Figure S2

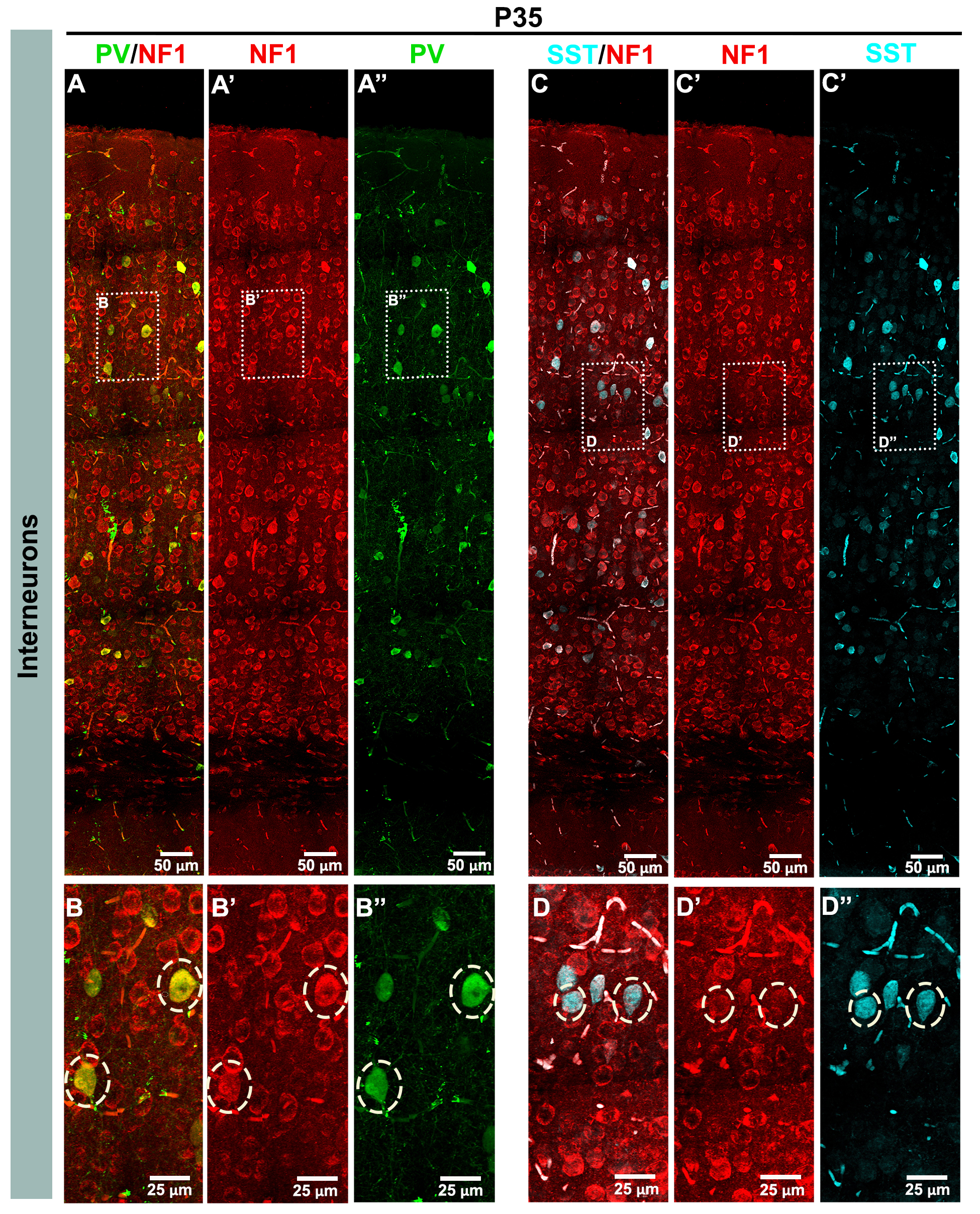

### Supplementary Figure S3

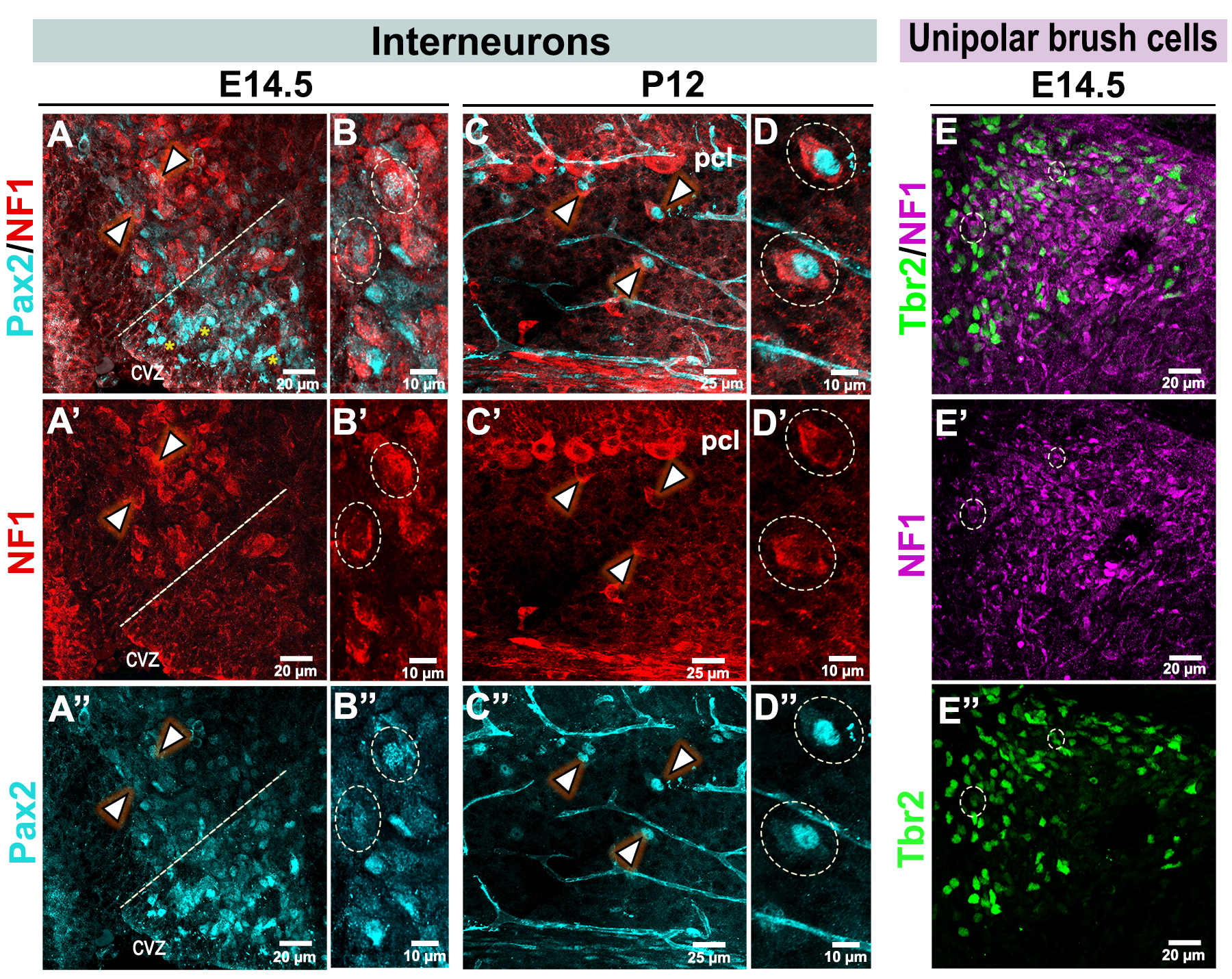
